# Fluoxetine treatment prevents cardiovascular changes in baroreflex and chemoreflex in rats subjected to chronic stress

**DOI:** 10.1101/2020.03.04.977033

**Authors:** Egidi Mayara Silva Firmino, Luciana Bärg Kuntze, Davi Campos Lagatta, Daniel Penteado Martins Dias, Leonardo Barbosa Moraes Resstel

**Affiliations:** Department of Pharmacology, School of Medicine of Ribeirão Preto, University of São Paulo, Ribeirão Preto, SP, 14090-090, Brazil; Barão de Mauá University Center, Ribeirão Preto, SP, 14090-180, Brazil

**Author notes:** **Corresponding author:** Leonardo Barbosa Moraes Resstel, Department of Pharmacology, School of Medicine of Ribeirão Preto, University of São Paulo, Ribeirão Preto, SP, 14090-090, Brazil. **Summary statement:** Our results provide evidence that autonomic changes may reflect important mechanisms in the etiology of cardiovascular diseases associated with exposure to chronic stress and fluoxetine treatment prevents these changes cardiovascular.

**Keywords:** Chronic stress, Baroreflex, Chemoreflex, Fluoxetine

## Abstract

Stress may influence the autonomic regulation, pathogenesis of cardiovascular disease and play an important role in animal behavior as depression. Depression is evidenced as a significant risk factor for cardiovascular changes. This is of great importance as some studies show an association between symptoms of depression and increased risk of cardiovascular disease (CVD) morbidity. Additionality, those alterations can be alleviated by use of antidepressants such as fluoxetine, which is a selective serotonin reuptake inhibitor (SSRIs). The link between depression and cardiovascular changes is known to be mediated in part by autonomic mechanisms that contribute to the regulation of cardiovascular function. However, studies on the effects of SSRIs on cardiovascular autonomic function are inconsistent. Thus, in the present study we investigated, in adult male rats, the effect of chronic and acute treatment with fluoxetine on changes in autonomic mechanisms of baroreflex and chemoreflex induced by the repeated restraint stress (RRS) or chronic variable stress (CVS) on baroreflex and chemoreflex in a protocol of 14 days of stress sessions. The results found demonstrated that exposure to chronic stress (RRS and CVS) promove changes on cardiovascular and ventilatory responses controlled by autonomic reflexes, such as baroreflex and chemoreflex. Additionality, that chronic fluoxetine treatment for 21 days was able to prevent not only anhedonic behavior, but also of autonomic changes cardiovascular induced by chronic stress. Taken together, our results show that pharmacological treatment with fluoxetine may be also helpful to prevent cardiovascular events on account of depressive states, by correcting alterations in autonomic function.

## 1. Introduction

Chronic stress exposure induces alterations of cardiovascular parameters such as blood pressure (BP) and heart rate (HR) (Crestani, 2016; Nakata et al., 1993). Furthermore, some studies suggest that stress may influence the autonomic regulation and pathogenesis of cardiovascular disease (Crestani, 2016; Firmino et al., 2019; Grippo et al., 2002, 2008). In addition to cardiovascular changes, chronic stress can also play an important role in altering animal behavior, known as anhedonia, similar to depression in humans (Willner et al., 1992). Depression is evidenced as a significant risk factor for cardiovascular changes (Low et al., 2010). This is of great importance as some studies show an association between symptoms of depression and increased risk of cardiovascular disease (CVD) morbidity (Bahall, 2019; Ivanovs et al., 2018). It is known that most animal models of depression are induced by uncontrollable exposure to stress. In this sense, several studies in the literature report this exposure to stress generates cardiovascular, in addition to depressive behaviors, that can be alleviated by use of antidepressants such as fluoxetine, a selective serotonin reuptake inhibitor (SSRIs) (Alboni et al., 2017; Ampuero et al., 2015; Grippo et al., 2006; Lu et al., 2017; Willner, 1990).

SSRIs are widely prescribed for treatment of depression (Jakubovski et al., 2016). In addition to having an antidepressant effect, SSRIs can also modify cardiovascular function. McFarlane, et al. (2001) demonstrated that SSRIs seems to facilitate the recovery of cardiac autonomic function in depressed patients (McFarlane et al., 2001). Additionally, chronic fluoxetine treatment caused mild hypertension, tachycardia and abnormality in the baroreflex function (Hong et al., 2017) that is a neuronal mechanism responsible for maintaining the BP at homeostatic levels by influencing variables, such as HR. However, some evidence suggests that SSRIs may also be associated with cardiovascular side effects (Pacher and Kecskemeti, 2004). Although some studies have investigated the effect of chronic fluoxetine treatment on cardiovascular stress responses, the results are contradictory (Grippo et al., 2006; Roche et al., 2007).

A recent study from our group showed that chronic stress promotes alterations not only in depressive behavior, but also in autonomic mechanisms of baroreflex, chemoreflex and cardiocirculatory variability (Firmino et al., 2019). In addition to baroreflex, the chemoreflex is another relevant neural mechanism responsible to evoke respiratory and cardiovascular responses due to specific hypoxic or hypercapnia conditions (Fitzgerald, 2000). This way, the mechanisms of baroreflex and chemoreflex play a major role in control of cardiovascular system through profound influences on autonomic outflow. The link between depression and cardiovascular changes is known to be mediated in part by autonomic mechanisms that contribute to the regulation of cardiovascular function (Grippo et al., 2008). Therefore, alterations caused by chronic stress seem to constitute important mechanisms in etiology related to cardiovascular disease (Firmino et al., 2019). However, studies on the effects of SSRIs on cardiovascular autonomic function are inconsistent.

Thus, the present study was designed to investigate the effect of chronic and acute treatment with fluoxetine on changes in autonomic mechanisms of baroreflex and chemoreflex induced by chronic stress.

## 2. Methods

### Animals

A total of 110 male Wistar rats aging 60 days-old (adult) were used in the present study. All animals were kept in the Animal Care Unit of the School of Medicine of Ribeirão Preto, University of São Paulo Ribeirão Preto (USP). The animals were housed in groups of five per cage in a temperature-controlled room (24±1°C) under a 12h light/dark cycle (lights on at 06:00 A.M.) and were given *water* and *food* ad *libitum. All* experiments were performed in accordance with Ethical Principles for Animal Experimentation followed by the Brazilian Committee for Animal Experimentation (COBEA) and approved by the Committee of Ethics in Animal Research of the School of Medicine of Ribeirão Preto, University of São Paulo *(number* 175/2014).

### Chronic Stress Regimens

The chronic stress regimens were based in protocols previously used and modified accordingly (Cruz et al., 2012; Duarte et al., 2015; Marin et al., 2007). Two animal models of stress were chosen. The repeated restraint stress (RRS) was used as a homotypic stressor, whereas chronic variable stress (CVS) was used as heterotypic stress stimuli. The animals of RRS group were restrained in plastic cylindrical tubes, 17 cm long and 7.5 cm high for 1 hour daily starting at 10:00 AM for 14 consecutive days. The CVS protocol consisted on exposure to different stressors in variable schedules for 14 consecutive days (Table 1). The protocols were initiated seven days apart from each other, so that the experiments performed with the respective groups also complied with this interval. Animals of the control groups were left undisturbed, except for cleaning the cages. All test described below were made in the same animals except the behavioral.

**TABLE 1.** Protocol of CVS

| *Day* | *Stress Type and Schedule* |
| --- | --- |
| *1* | 10:00 AM restraint stress, 60 min; 7:00 PM, humid sawdust, overnight |
| *2* | 3:00 AM, cold (4°C) isolation, 60 min; 7:00 PM, lights in, overnight |
| *3* | 12:00 AM, lights off, 180 min; 3:00 PM, swim stress, 4 min |
| *4* | 07:30 AM, humid sawdust, all day; 7:00 PM, food/water deprivation, overnight |
| *5* | 1:00 PM, swim stress, 3 min; 7:00 PM, isolation housing overnight |
| *6* | 2:00 PM, cold (4°C) isolation, 15 min; 3:00 PM, lights off, 120 min |
| *7* | 7:00 PM, humid sawdust and lights on, overnight |
| *8* | 7:00 PM, isolation and food/water deprivation, overnight |
| *9* | 4:00 PM, restraint stress, 60 min; 7:00 PM, lights on, overnight |
| *10* | 09:00 AM, swim stress 4 min; 10:00 AM, restraint stress, 60 min |
| *11* | 3:00 AM, cold (4°C) isolation, 60 min; 7:00 PM, food/water deprivation, overnight |
| *12* | 10:00 AM restraint stress, 60 min; 7:00 PM, humid sawdust, overnight |
| *13* | 07:30 AM, humid sawdust, all day; 7:00 PM, food/water deprivation, overnight |
| *14* | 10:00 AM restraint stress, 60 min |

CVS = chronic variable stress.

**TABLE 2.** Sigmoidal curve parameters of animals submitted to the chronic stress protocol that received chronic or acute treatment with fluoxetine

| Groups | G<br>(bpm/mmHg) | P1<br>(bpm) | P2<br>(bpm) | $\Delta P$<br>(bpm) | PA<sub>50</sub><br>(mmHg) |
| --- | --- | --- | --- | --- | --- |
| **Control + Veh** | -1.64 ± 0.03 | -84.4 ± 2,7 | 75.14 ± 3 | 159 ± 9.09 | 3.01 ± 2.67 |
| **Control + Chronic Flx** | -2.38 ± 0.05<sup>#</sup> | 53.8 ± 3.3<sup>#</sup> | 48.8 ± 3.5<sup>#</sup> | 102.7 ± 15<sup>#</sup> | -3.52 ± 4.01 |
| **Control + Acute Flx** | 1,85 ± 0,064 | -69,6 ± 2,5<sup>#</sup> | 77,8 ± 5 | 147,4 ± 14 | -1,59 ± 2,22 |
| **RRS + Veh** | -1.09 ± 0.05<sup>*</sup> | -113.7 ± 4.1<sup>*</sup> | 102.3 ± 1.0<sup>*</sup> | 216 ± 9.2<sup>*</sup> | 1.66 ± 2.43 |
| **RRS + Chronic Flx** | -1.91 ± 0.04<sup>#</sup> | -86 ± 2.7<sup>#</sup> | 70.3 ± 1.4<sup>#</sup> | 154 ± 11.4<sup>#</sup> | 3.05 ± 2.88 |
| **RRS + Acute Flx** | -1,23 ± 0,07<sup>#</sup> | -79,7 ± 0,9<sup>#</sup> | 94,2 ± 5,5 | 174 ± 14,2<sup>#</sup> | -0,82 ± 3,53 |
| **CVS + Veh** | -1.64 ± 0.03 | -83 ± 3.0 | 80.5 ± 4.1 | 170 ± 13.8 | 5.68 ± 4.44 |
| **CVS + Chronic Flx** | -1.39 ± 0.03 | -95 ± 8.0 | 79.6 ± 2.2 | 157 ± 11.6 | 1.28 ± 1.65 |
| **CVS + Acute Flx** | -1,71 ± 0,02 | -95 ± 8 | 75 ± 2,7 | 155 ± 9,6 | 2,67 ± 3,95 |
| **Condition:** | F<sub>(2,55)</sub>= 11.54 | F<sub>(2,55)</sub>= 6.31 | F<sub>(2,55)</sub>= 5.38 | F<sub>(2,55)</sub>= 45.15 | F<sub>(2,55)</sub>= 0.98 |
| **Treatment:** | F<sub>(2,55)</sub>= 10.42 | F<sub>(2,55)</sub>= 6.11 | F<sub>(2,55)</sub>= 3.55 | F<sub>(2,55)</sub>= 41.94 | F<sub>(2,55)</sub>= 1.01 |
| **Interaction:** | F<sub>(4,55)</sub>= 6.65 | F<sub>(4,55)</sub>= 1.17 | F<sub>(4,55)</sub>= 0.71 | F<sub>(4,55)</sub>= 5.95 | F<sub>(4,55)</sub>= 0.45 |

Values are mean ± SEM. \*P<0.05 vs control, # P<0.05 vs versus respective group vehicle within same condition. Two-way ANOVA followed by Bonferroni's *post hoc* test.

### Pharmacological treatment

The animals were randomly assigned into nine groups treated with either fluoxetine (10 kg/ml/kg, i.p.; Sigma-Aldrich, St. Louis, MO); or saline (0.9% saline solution + Tween-80, 1 ml / kg). The groups were divided as follows: 1) control/vehicle; 2) RRS/vehicle and 3) CVS/vehicle, received daily intraperitoneal (i.p.) injections of vehicle for 21 days; 4) control/chronic fluoxetine; 5) RRS/chronic fluoxetine and 6) CVS/chronic fluoxetine, received daily intraperitoneal (i.p.) injections of fluoxetine for 21 days; 7) control/acute fluoxetine; 8) RRS/acute fluoxetine and 9) CVS/acute fluoxetine, received daily intraperitoneal (i.p.), administration of vehicle 0.9% saline solution + Tween-80, 1 ml/kg) for 20 days and on the last day fluoxetine (10 mg / kg) was administered. All animals were treated for 21 days starting 7 days prior to stress protocols.

### Experimental protocols

All Animals were treated for 21 days starting and only animals of CVS and RRS groups were submitted to daily sessions of stress for 14 days. All chronic treatments started 7 days prior to stress protocols. On the 14th day, after the last session of stress, animals in all experimental groups were subjected to surgical preparation. Twenty-four hours later, tests were performed. On test days, animals were transferred to the experimental room in their home box and allowed 30 minutes to adapt to experimental room conditions before starting experiments. Subsequently, a 10 minutes period of basal cardiovascular activity was recorded. Afterwards, we performed baroreflex and chemoreflex stimulation following the methods mentioned above.

### Sucrose preference test

This test was developed with an independent group of animals. At the end of the chronic stress protocol, it was evaluated ingestion of sucrose (2% solution) as a measurement of anhedonia (decreased or absent ability to experience pleasure). For this purpose, the protocol was carried out in two days. In each day, the animals were separated individually. On the first day, animals were habituated with samples (sucrose solution or drinking water) for a period of 5 hours. In the following day (testing day), the initial bottles with pure water and sucrose solution were weighed according to protocol (initial weight) *(Willner, 1997)*. Then, the two bottles were offered to rats for a period of 10 hours. Thereafter, they were weighed again (final weight). Consumption of both water and sucrose solution was calculated as the difference between initial and final weights. The preference for sucrose (depicted in the formula below) was calculated as a percentage value. Sucrose consumption (initial weight minus the final weight in grams of sucrose bottles) and total consumption, which is water consumption (initial weight minus final weight in grams of water bottles) + sucrose consumption (initial weight minus the final weight in grams of sucrose bottles) were devided and multiplied by 100.

### Surgical procedures

Animals were anesthetized with tribromoethanol (250 mg/kg, i.p.) and a catheter (a 4 cm segment of PE-10 that was heat-bound to a 13 cm segment of PE-50 (Clay Adams, Parsippany, New Jersey, USA) was inserted into the abdominal aorta through the femoral artery, for blood pressure recording. A second catheter was implanted into the femoral vein for vasoactive drug infusion. Both catheters were tunneled under the skin and exteriorized on the animal’s dorsum. After the surgery, the animals were given polyantibiotic preparation of streptomycin and penicillin (0,3 ml, I.M; Pentabiotico; combination of penicillins and streptomycins; 80.000 UI; Fort Dodge, Campinas, SP,Brazil) to prevent infection and also received the non-steroidal anti-inflammatory flunixine meglumine (2.5 mg/Kg, S.C, Banamine; Intervet Schring-Plough, Cruzeiro, SP, Brazil) for postoperative.

### Measurement of cardiovascular parameters

The pulsatile arterial pressure of freely moving animals was recorded using an ML870 preamplifier (LabChart, USA) and an acquisition board (PowerLab, AD Instruments, USA) connected to a computer. Mean arterial pressure (MAP) and heart rate (HR) values were derived from pulsatile recordings and processed on-line.

### Baroreflex assessment

The baroreflex was activated by phenylephrine (α1 adrenoceptor agonist; 50 μg kg^−1^; 0.34 mL min^−1^) or SNP (NO donor; 50 μg kg^−1^; 0.8 mL min^−1^) infusion using an infusion pump (KD Scientific, Holliston, MA, USA). The phenylephrine or SNP infusion lasted 30–40s and caused, respectively, an increase and decrease in BP (Alves et al., 2009). Baroreflex curves were constructed, matching MAP variations with HR responses. Paired values for variations in MAP (ΔMAP) and HR (ΔHR) were plotted to create sigmoid curves for each rat, which were used to determine baroreflex activity (Resstel et al., 2004). To analyze bradycardic and tachycardic responses separately, HR values matching 10, 20, 30 and 40 mmHg of MAP changes were calculated (Alves et al., 2009). Values were plotted to create linear regression curves for each rat, and their slopes were compared to determine changes in baroreflex gain.

### Chemoreflex activation

The chemoreflex was activated by systemic bolus intravenous injections of 0.1 mL of KCN (40 μg per rat; Merck, Darmstadt, Germany) following procedures described by Franchini & Krieger (1993) and experimental conditions previously validated (Granjeiro et al., 2011). The magnitude of changes in MAP and HR in response to chemoreflex activation was quantified at the peak, and the duration of the responses was recorded (Granjeiro et al., 2011; Kuntze et al., 2016).

The measurements of respiratory frequency response (*f*_R_) induced by chemoreflex activation were evaluated by whole body plethysmography method for small animals described by (Bartlett and Tenney, 1970). Rats were kept inside a 6-L Plexigas® chamber (with sealed exit ports for catheters) which allowed them to move freely. After the chamber was closed to *f*_R_ recording, pressure oscillations originated by breathing movements were detected by a differential pressure transducer (ML141 Spirometer, PowerLab, ADInstruments, Bella Vista, NSW, Australia). The *f*_R_ was expressed as number of cycles per minute (cpm), which was calculated from the extrapolation of the plestimographic signal, regarding the volume calibration accomplished for each measurement (1 ml injection of air into the chamber). Respiratory cycles were acquired during 20 s before and 20 s after chemoreflex activation with KCN (I.V.) and was analyzed into 2 s intervals by acquisition software program (PowerLab, ADInstruments, Bella Vista, NSW, Australia) with manual corrections, when necessary. Equations described by Drorbaugh e Fenn (1955) allowed calculation of the tidal volume (*V*_T_) and the minute ventilation (*V*_E_) was obtained as the product of *f*_R_ and *V*_T_. The *V*_T_ and *V*_E_ parameters were only calculated 10 points before chemoreflex activation to avoid wrong values given that vigorous behavioral responses are evoked by KCN injection.

### Data Analysis

All the data present a normal distribution and homogeneity of variance. Data were expressed as mean ± SEM. Results were analyzed using two-way analysis of variance ANOVA following the Bonferroni’s post hoc test for identification of differences between the groups the from GraphPad Prism Software version 7 (San Diego, CA, USA). Results of statistical tests with *p<0.05* were considered significant.

## 3. Results

### 3.1 Effect of chronic stress and fluoxetine treatment on basal cardiovascular parameters and sucrose preference test

#### Sucrose preference test

Analysis of sucrose preference indicated a main effect of stress (F_2,24_ = 6.43, *p*<*0.05)*, as well of fluoxetine treatment (F_2,24_ = 15.24, *p*<*0.05)*. However, there was no interaction between the factors (F_2,24_ = 2.715, *p*<*0.05)*. *Post-hoc* analysis revealed that exposure to repeated restrain or variable stress reduced the sucrose preference as compared to group control (*p*<*0.05*). In addition, fluoxetine treatment prevents these changes caused by stress (*p*<*0.05*) (Fig.1).

**Figure 1.**
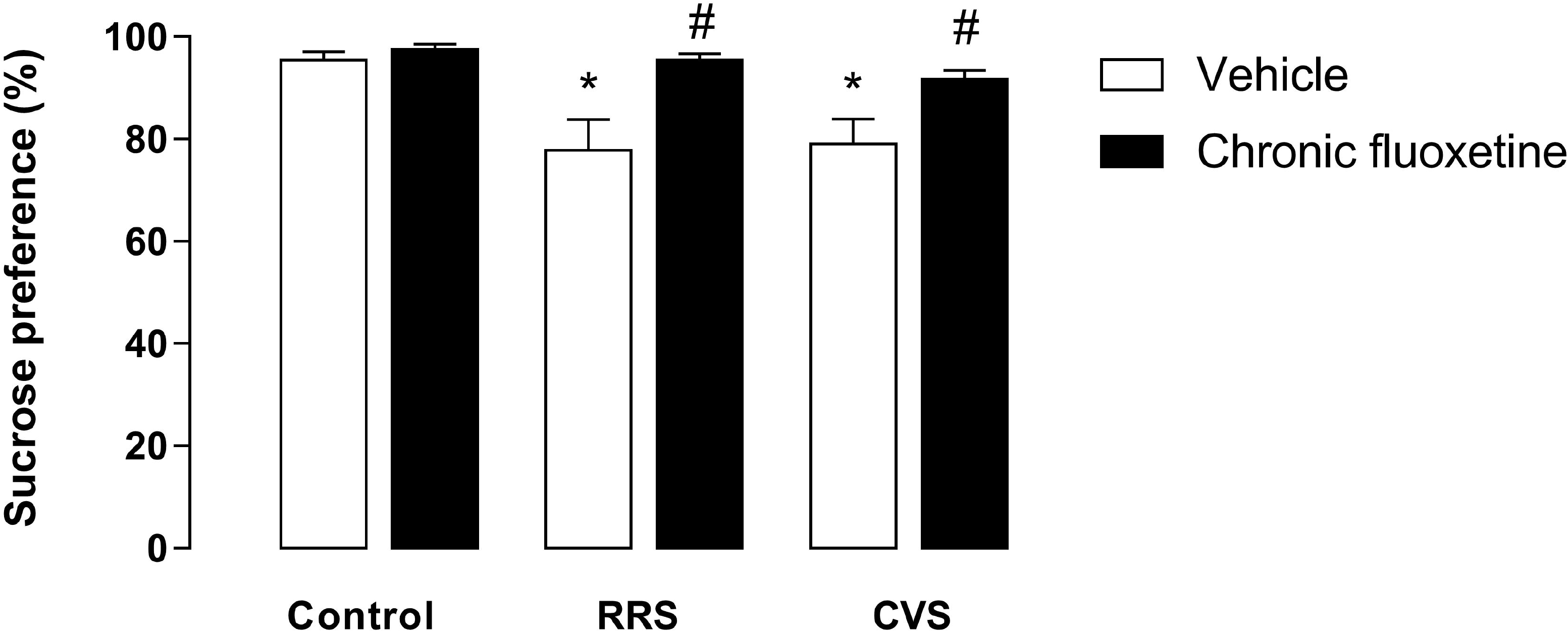
Effect of chronic fluoxetine treatment on the sucrose preference test in control and stressed rats (RRS or CVS). (A) Consumption of 2% sucrose solution, *P<0.05 versus control group, One-way ANOVA. *P<0.05 versus control group, Two-way- ANOVA. The bars represent the mean ± SEM of the control (n=5), repeated chronic stress (RRS, n=5) and chronic variable stress (CVS, n=5) followed by Bonferroni’s *post hoc* test. ANOVA = analysis of variance.

#### Basal cardiovascular parameters

The protocols of chronic stress did not affected mean arterial pressure (MAP) (F_2,71_ = 0.4183; *p>0.05*) an HR (F_2,71_ = 2.075; *p>0.05*). The chronic or acute fluoxetine treatment too did not affected MAP (F_2,71_ = 0.765; *p>0.05*) and HR (F_2,71_ = 1.222; *p>0.05*); the ANOVA yielded no significant main effects or interaction >0.05 for all comparisons (Fig.2).

**Figure 2.**
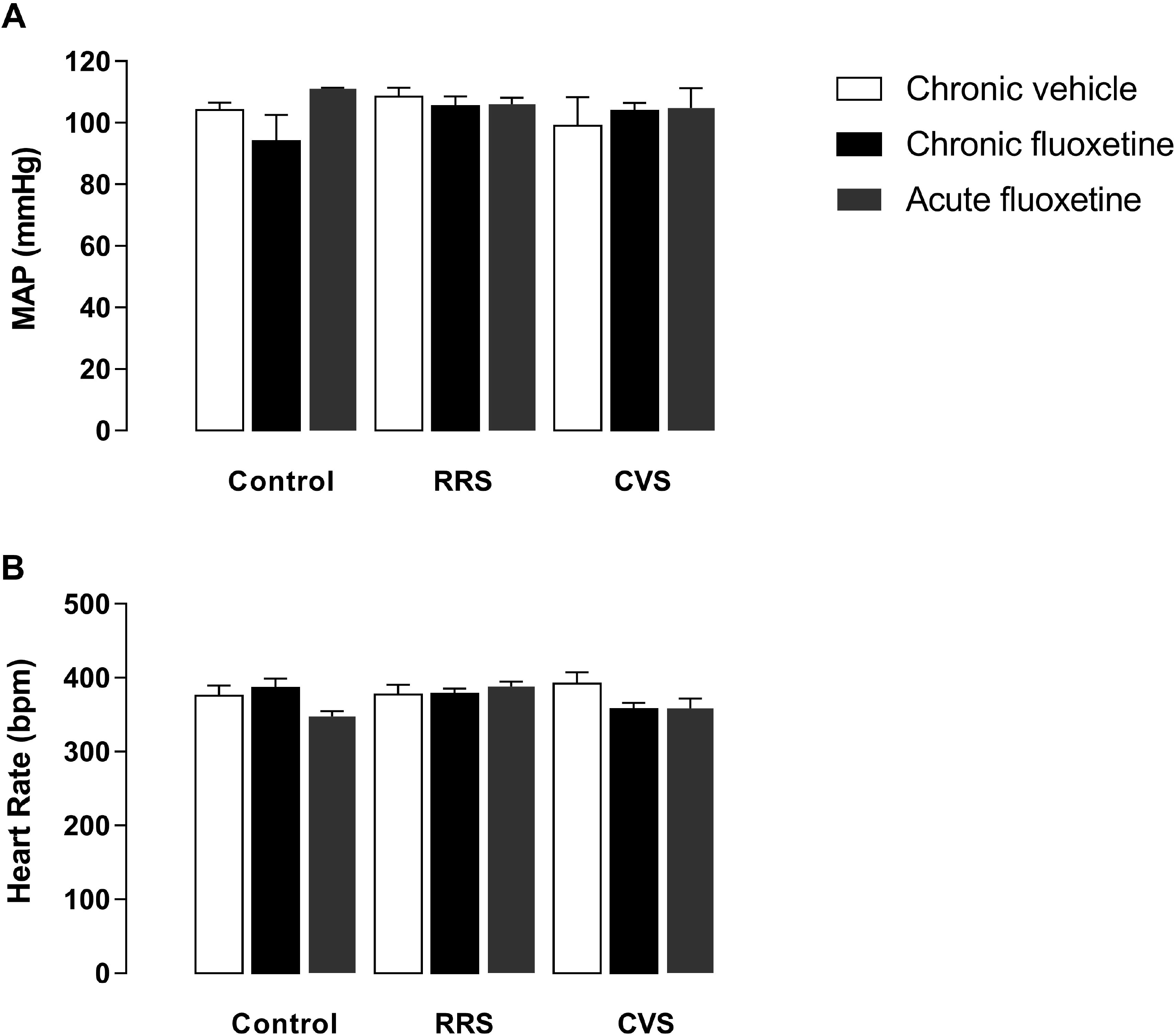
Effect of chronic or acute fluoxetine treatment on **(A)** Baseline mean arterial pressure (MAP) and **(B)** baseline heart rate (HR) in control, RRS and CVS rats. The bars represent mean ± SEM of the control (Control: vehicle (n=8); chronic fluoxetine (n=11) and acute fluoxetine (n=6)); repeated chronic stress (RRS: vehicle, n=9; chronic fluoxetine (n=11) and acute fluoxetine (n=8) and chronic variable stress (CVS: vehicle n=11; chronic fluoxetine (n=10) and acute fluoxetine (n=6)). *P<0.05 was considered statistically different. ANOVA two-way, followed by Bonferroni’s *post hoc* test. ANOVA = analysis of variance.

### 3.2 Effect of chronic stress and fluoxetine treatment on cardiac baroreflex activity

Nonlinear regression analysis of baroreflex activity indicated effect of stress (bradycardic: F_2,55_ = 10.33; *p<0.05;* tachycardic: F_2,55_ = 3.77; *p<0.05*) and treatment (bradycardic: F_2,55_ = 8.06; *p<0.05;* tachycardic: F_2,55_ = 2.21; *p<0.05*), but there was no interaction between condition x treatment (F_4,55_ = 1.26; *p>0.05*). *Post hoc* analysis revealed that the RRS/vehicle increased the slope of linear regression of the bradycardic (control/vehicle = −2.33 ± 0.23 and RRS/vehicle = −3.39 ± 0.35; *p<0.05*) and tachycardic responses (control/vehicle = −1.98 ± 0.30 and RRS/vehicle = −2.57 ± 0.24; *p<0.05*). However, there was no significant difference between the CVS/vehicle group and the control/vehicle group in bradycardic (control/vehicle = −2.33 ± 0.23 and CVS/vehicle = −2.56 ± 0.14) and tachycardic responses (control/vehicle = −1.59 ± 0.18 and CVS/vehicle = −2.15 ± 0.38). Chronic fluoxetine treatment shows an effect per se, as it decreases the slope of the bradycardic curve when compared to control/vehicle group (control/vehicle = −2.33 ± 0.23 and control/chronic fluoxetine = −1.29 ± 0.18; *p<0.05)*. On the other hand, it was able to prevent the bradycardic enhancement induced by repeated stress; (RRS/vehicle = −3.39 ± 0.35 and RRS/chronic fluoxetine = 2.35 ± 0.28; *p<0.05;* CVS/vehicle = 2.56 ± 0.14; CVS/chronic fluoxetine = 2 42 ± 0.29; *p>0.05*). The same was observed for the tachycardic component of the baroreflex (control/vehicle = −1.98 ± 0.30 VS control/chronic fluoxetine = −1.15 ± 0.23; *p<0.05;* RRS/vehicle = −2.57 ± 0.24 VS RRS/chronic fluoxetine −1.91 ± 0.22; *p<0.05*; CVS/vehicle = −2.15 ± 0.38 VS CVS/chronic fluoxetine −1.93 ± 0.30; *p>0.05*) (Fig. 3). Additionally, acute fluoxetine treatment did not alter the tachycardic component of the baroreflex in either group *(p>0.05)*, but reduces the bradycardic component of the RRS group (RRS/vehicle = −3.396 ± 0.35; RRS/acute fluoxetine = −2.27 ± 0.12; *p<0.05*) (Fig.4). Apart from BP50, the sigmoid curve parameters (P1, P2, G, ΔP) were increased in the RRS group (Table 1).

**Figure 3.**
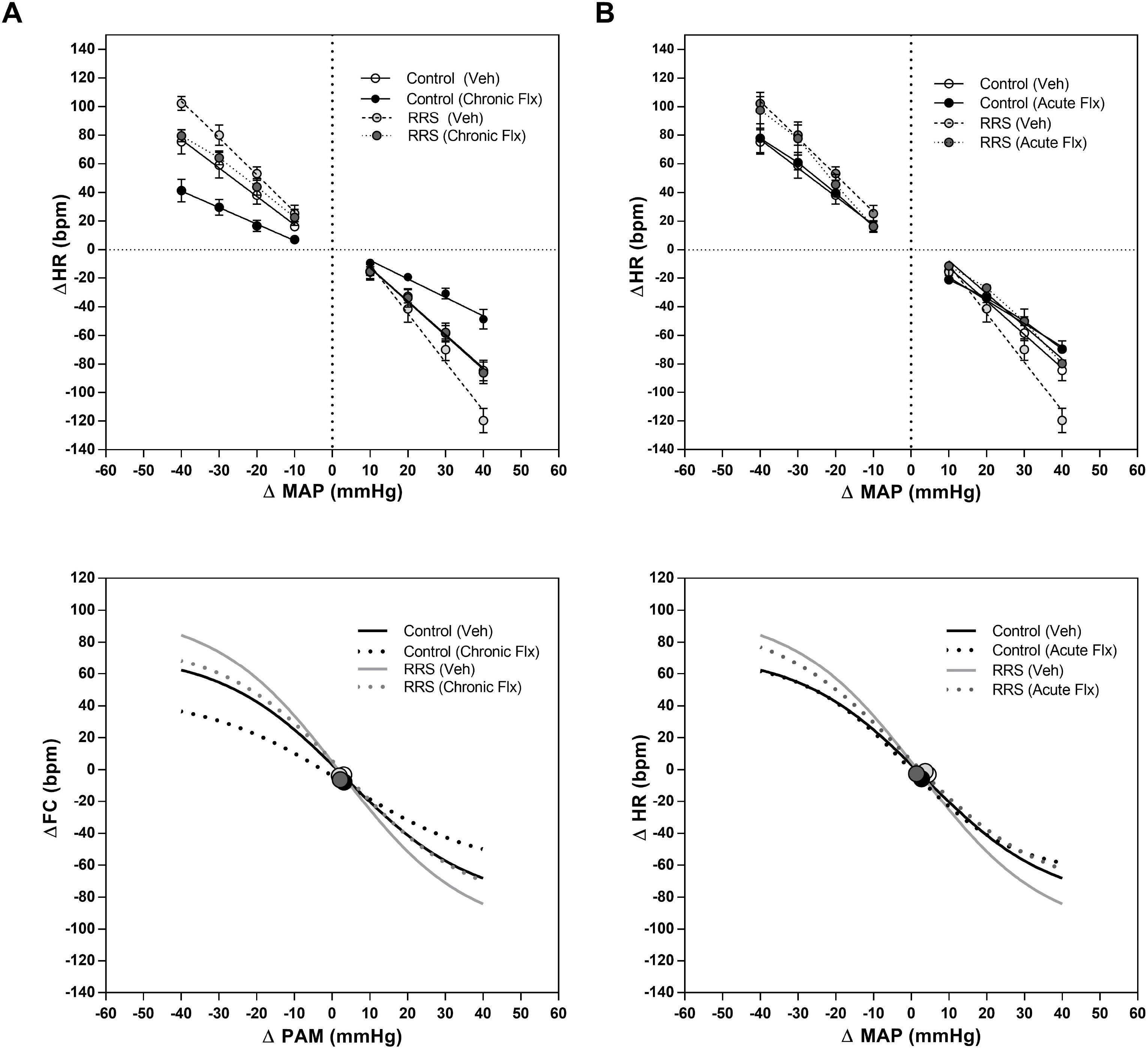
**(A)** Linear regression curves (top panel) and sigmoid curves (bottom panel) correlating the ΔMAP and ΔHR responses of control (Control: vehicle (n=7); chronic fluoxetine (n=8)) and repeated chronic stress (RRS: vehicle, (n=8); chronic fluoxetine (n=8)). **(B)** Linear regression curves (top panel) and sigmoid curves (bottom panel) correlating the ΔMAP and ΔHR responses of the control (Control: vehicle (n=7); chronic fluoxetine (n=8)) and chronic variable stress (CVS: vehicle (n=9); chronic fluoxetine (n=8)). Values are means ± SEM. bpm, beats min^−1^. The circles in the sigmoidal curves represent the BP50.

**Figure 4.**
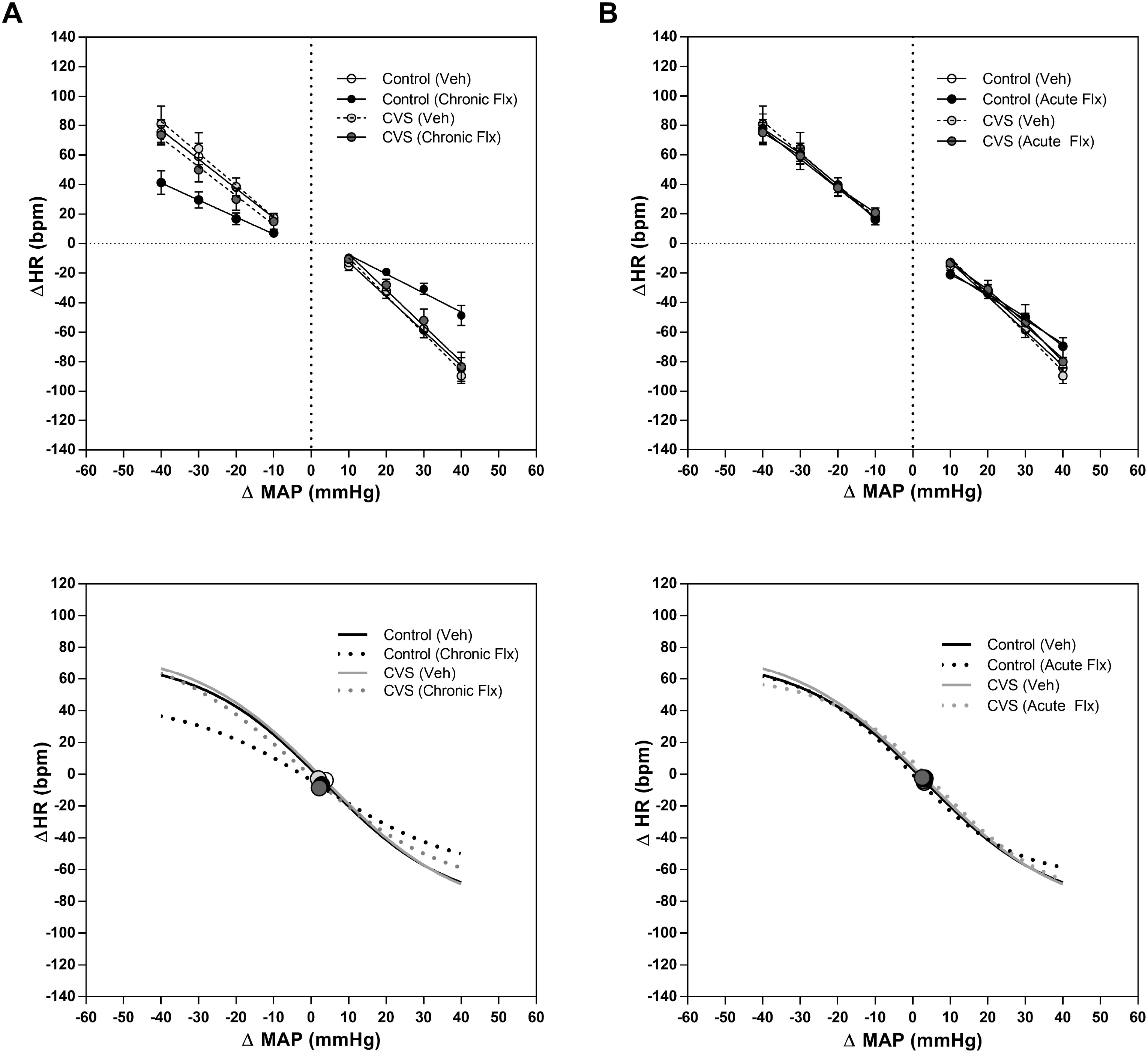
**(A)** Linear regression curves (top panel) and sigmoid curves (bottom panel) correlating the ΔMAP and ΔHR responses of control (Control: vehicle (n=7); acute fluoxetine (n=5)) and repeated chronic stress (RRS: vehicle, (n=8); acute fluoxetine (n=8). **(B)** Linear regression curves (top panel) and sigmoid curves (bottom panel) correlating the ΔMAP and ΔHR responses of the control (Control: vehicle (n=7); chronic; acute fluoxetine (n=5)) and chronic variable stress (CVS: vehicle n=9; acute fluoxetine (n=6)). Values are means ± SEM. bpm, beats min^−1^. The circles in the sigmoidal curves represent the BP50.

### 3.3 Effect of chronic stress and fluoxetine treatment on cardiovascular and respiratory frequency to chemoreflex activation and on baseline respiratory parameters

#### Cardiovascular responses to chemoreflex activation

The data summarized in Fig. 5A show effect of stress (F_2,69_ = 17.15; *p<0.05*), treatment (F_2,69_ = 35.67; *p<0.05*), and stress x treatment interaction (F_4,69_ = 17,53; *p<0.05*) in the magnitude of the pressor response to chemoreflex activation in rats. *Post hoc* analysis revealed that the RRS/vehicle and CVS/vehicle groups decreased the magnitude of the pressor response to chemoreflex activation when compared to the control/vehicle group (*p<0.05*) (Fig. 5A). Chronic fluoxetine treatment promotes a decreased pressor response in comparison with control/vehicle group, whereas chronic and acute fluoxetine treatment were able to increase the magnitude of the pressor response when compared to their respective vehicles (*p<0.05*). The magnitude of the bradycardic response to chemoreflex activation there was no significant difference between groups (F_2,69_ = 0.605, *p>0.05*) (Fig. 5B). There was no effect of chronic fluoxetine treatment (*p>0.05*), but acute fluoxetine treatment was able to reduce the magnitude of the bradycardic response in the groups submitted to chronic stress protocol (F_2,69_ = 8.378, *p<0.05*).

#### Respiratory frequency to chemoreflex activation

KCN-induced chemoreflex activation demonstrated effect of stress (F_2,69_ = 6.26, *p<0.05)*, and treatment (F_2,69_ = 7.34, *p<0.05*), but there is no interaction between stress and treatment (F_4,69_ = 1.50, *p>0.05*) in respiratory frequency. *Post hoc* analysis revealed that the RRS/vehicle and CVS/vehicle groups increased respiratory frequency when compared to control/vehicle (*p>0.05*). However chronic and acute fluoxetine treatment further potentiation respiratory frequency in the control and RRS (*p>0.05*) group but not in the CVS (*p<0.05*) (Fig.6).

#### Baseline respiratory parameters

Regarding baseline ventilatory parameters, results showed an effect of stress (F_2,69_ = 3.646, *p<0.05*), treatment (F_2,69_ = 5.743, *p<0.05*) and stress x treatment interaction (F_4,69_ = 6.725, *p<0.05)* in baseline respiratory frequency values. It is possible to observe that the CVS/vehicle group had a higher baseline respiratory rate compared to the control/vehicle group, which was decreased after chronic fluoxetine treatment (*p<0.05*). Acute treatment promoted an increase in respiratory rate only in the control/vehicle group, with no effect on the other groups (*p>0.05)* (Fig 7A). Additionally, both protocols of chronic stress promoted an attenuation in the tidal volume responses (*V*_T_); (F_2,69_ = 6.076, *p<0.05*) and minute ventilation (*V*_E_); (F_2,69_ = 7.907, *p<0.05*). However, only chronic fluoxetine treatment presented an effect in the control group on both baseline ventilatory parameters (*p<0.05*) (Fig. 7B and C).

## 4. Discussion

The present study reinforced the effect of different types of chronic stress (RRS and CVS) on cardiovascular and ventilatory responses controlled by autonomic reflexes, such as baroreflex and chemoreflex, as previously shown by our group(Firmino et al., 2019). On the other hand, the great novelty of the present study is that chronic fluoxetine treatment for 21 days was able to prevent not only anhedonic behavior, but also most part of autonomic changes cardiovascular induced by chronic stress.

The preference for sucrose has been used as a marker of chronic stress exposure in over 50 studies, indicating a depressive-like behavior in animals (Willner, 2005). In this way, several studies report that chronic exposure to stress generates behaviors modifications, that can be alleviated by the use of antidepressants such as fluoxetine, which is a SSRIs (Alboni et al., 2017; Ampuero et al., 2015; Grippo et al., 2006; Lu et al., 2017; Willner, 1990).

In addition to behavior changes, chronic stress can also alter cardiovascular function. Several studies have shown that chronic exposure to stress can lead to increased BP and HR, depending on the type of stress protcol (Barres et al., 2004; Carrive, 2006; Duarte et al., 2015; Grippo et al., 2008; Irvine et al., 1997; Tavares and Correa, 2006). However, our results did not show any effect of chronic stress on such parameters. On the other hand, our findings are consistent with studies by Grippo et al (2006) (Grippo et al., 2006).and Firmino et al (2019)(Firmino et al., 2019) who demonstrated that chronic stress was unable to alter baseline BP and HR. Moreover, treatment with fluoxetine did not influence these parameters either. These results are in agreement with studies performed by Almeida et al. (2015) (Almeida et al., 2015) and Roose et al. (1998) (Roose et al., 1998) in which chronic stress and SSRI treatment do not modify basal parameters.

However, our data corroborate a previous study by our group showing that RRS but not CVS was able to potentiate tachycardic and bradycardic responses to baroreflex activation induced by vasoactive drugs. An additional protocol included in the present study, demonstrated that fluoxetine treatment for a period of 21 days normalized tachycardia and bradycardia magnitude that had been enhanced by RRS. In addition, reduction in both plateaus of sigmoid curves suggested that chronic fluoxetine treatment led to this effect by decreasing both sympathetic and parasympathetic outflows. In the study of Crestani et al (2011) it was shown that chronic fluoxetine treatment for 21 days reduces sympathetic component of baroreflex activity and facilitates the parasympathetic component of baroreflex activity, but these results were obtained with acute stress (Crestani et al., 2011). Additionally, our results demonstrated chronic fluoxetine treatment decreased the tachycardia and bradycardia reflex responses of the control group. This can be considered a collateral effect of the treatment. Hong et al (2017) shown that fluoxetine treatment for 14 weeks is capable of promoting abnormality in the baroreflex function which might be attributed sympathoexcitation and probably parasympathetic depression in conscious normal rats (Hong et al., 2017). Regarding acute fluoxetine treatment, our results show that acute treatment did not interfere with tachycardic response, however it was able to attenuate the bradycardic response of the RRS group, analyzed by linear regression curves.

Some mechanisms may explain the effect of fluoxetine on baroreflex activity. In mammals, baroreceptors are present in nerve endings found in the carotid sinus and aortic arch (Papademetriou et al., 2011). However, structures of the limbic system and forebrain have been shown to connect to brainstem regions responsible for modulation of autonomic responses (Berntson, 2007). In this context, studies have shown that the stimulation or inhibition of these limbic structures, such as medial prefrontal cortex (Lagatta et al., 2015; Resstel et al., 2004; Resstel et al., 2006), hippocampus (Ferreira-Junior et al., 2017) is able to influence the function of the baroreflex. Moreover, chronic 21-day fluoxetine treatment increases Fos protein expression, a marker of neuronal activation, in those supramedullary areas involved in controlling autonomic activity (Lino-de-Oliveira et al., 2001). Accordingly, it is possible that central action of fluoxetine could mediate these alterations in baroreflex.

In addition to modifications in baroreflex activity, chronic stress also promoted deregulation in the activity of the chemoreflex. Our results demonstrate that RRS and CVS modified chemoreflex cardiovascular parameters, since both protocols reduced the magnitude of the pressor response without changing magnitude of bradycardia. Additionally, chronic treatment with fluoxetine prevented an increase in the magnitude of pressure response. However, acute treatment with fluoxetine was able to potentiate the pressor response in all groups and attenuated bradycardic response in RRS and CVS groups. According to Paton et al., (2001), activation of chemoreflex with KCN generates impulses that are conducted to the nucleus of the solitary tract (NTS) (Paton et al., 2001). NTS receives projections of serotonergic neurons from the caudal raphe nucleus (Thor and Helke, 1987) Thus, decreased serotonin in the dorsomedial bulb attenuates both ventilatory responses and sympathetic responses resulting from the administration of KCN (Kung and Scrogin, 2011). Therefore, serotonin in this region would facilitate the activity of chemoreflex, which could justify the increase in the magnitude of pressor response after treatment with fluoxetine. However, when activation of the obscure raphe nucleus occurs, it releases serotonin in NTS, playing an inhibitory role on the bradycardic response to hypoxia situation and consequent activation of chemoreflex (Weissheimer and Machado, 2007).It probably leads to a reduction in bradycardic response evoked by acute administration with fluoxetine.

In addition to cardiovascular changes, RRS and CVS also caused respiratory changes, by decreasing baseline *V*_E_ and *V*_T_ values and an increase in the magnitude of *f*_R_ in response to activation of chemoreflex. Additionally, chronic and acute treatment with fluoxetine potentiated *f*_R_ in response to activation of chemoreflex., while it did not change the baseline parameters of *V*_E_ and *V*_T_. The different ventilatory responses found after chronic and acute treatment with fluoxetine, probably occur due to different modulating effects of 5-HT on multiple receptors involved in respiratory control (Hodges and Richerson, 2008). Consequently, there are several different receptor subtypes 5-HT expressed in the respiratory nuclei (Lein et al., 2007) that may be mediating different responses. Some studies suggest that the bulbar raphe nuclei are not activated in baseline conditions, since the microinjection of fluoxetine or muscimol (GABAA receptor agonist) in the bulbar raphe did not cause changes in ventilation (Hodges et al., 2004; Taylor et al., 2004; Taylor et al., 2006), suggesting the participation of other nuclei in these responses. Thus, based on the fact that respiratory response to hypoxia is one of the most important components of activation of chemoreflex, it is concluded that all the mechanisms involved in the generation of respiratory activity are quite complex due to the different nuclei involved in this response, as raphe nuclei, NTS and ambiguous nucleus(Fuxe, 1965; Holtman, 1988; Li et al., 1993)

In this way, these results have provided evidence that the autonomic alterations could reflect important mechanisms underlining the etiology of cardiovascular diseases in depressive states, which are associated with chronic stress exposure. Additionality, the present study demonstrated that treatment with fluoxetine seems to preclude several cardiovascular alterations both in baroreflex and chemoreflex caused by chronic stress. Taken together, our results show that pharmacological treatment with fluoxetine may be also helpful to prevent cardiovascular events on account of depressive states, by correcting alterations in autonomic function.

## Abbreviations definition list

BP50 - medium blood pressure; CVD - cardiovascular disease; CVS - chronic variable stress; *f*_R_ - respiratory frequency; G - average gain; HR - heart rate; KCN - Potassium cyanide; MAP - mean arterial pressure; NTS - nucleus of the solitary tract; P1 - lower plateau; P2 - upper plateau; PA - pressure arterial; RRS - repeated restraint stress; SNP - sodium nitroprusside; SSRIs - selective serotonin reuptake inhibitor; *V*_E_ - minute ventilation; *V*_T_ - tidal volume; ΔP - heart rate range

## Acknowledgments

The authors wish to thank Camargo, L.H.A. and Mesquita O. for technical help.

## Author Contributions

EMSF, LBK, DCL and DPD contributed conception and design of the study; LMR organized the database and performed the statistical analysis. All authors contributed to manuscript revision, read and approved the submitted version

---

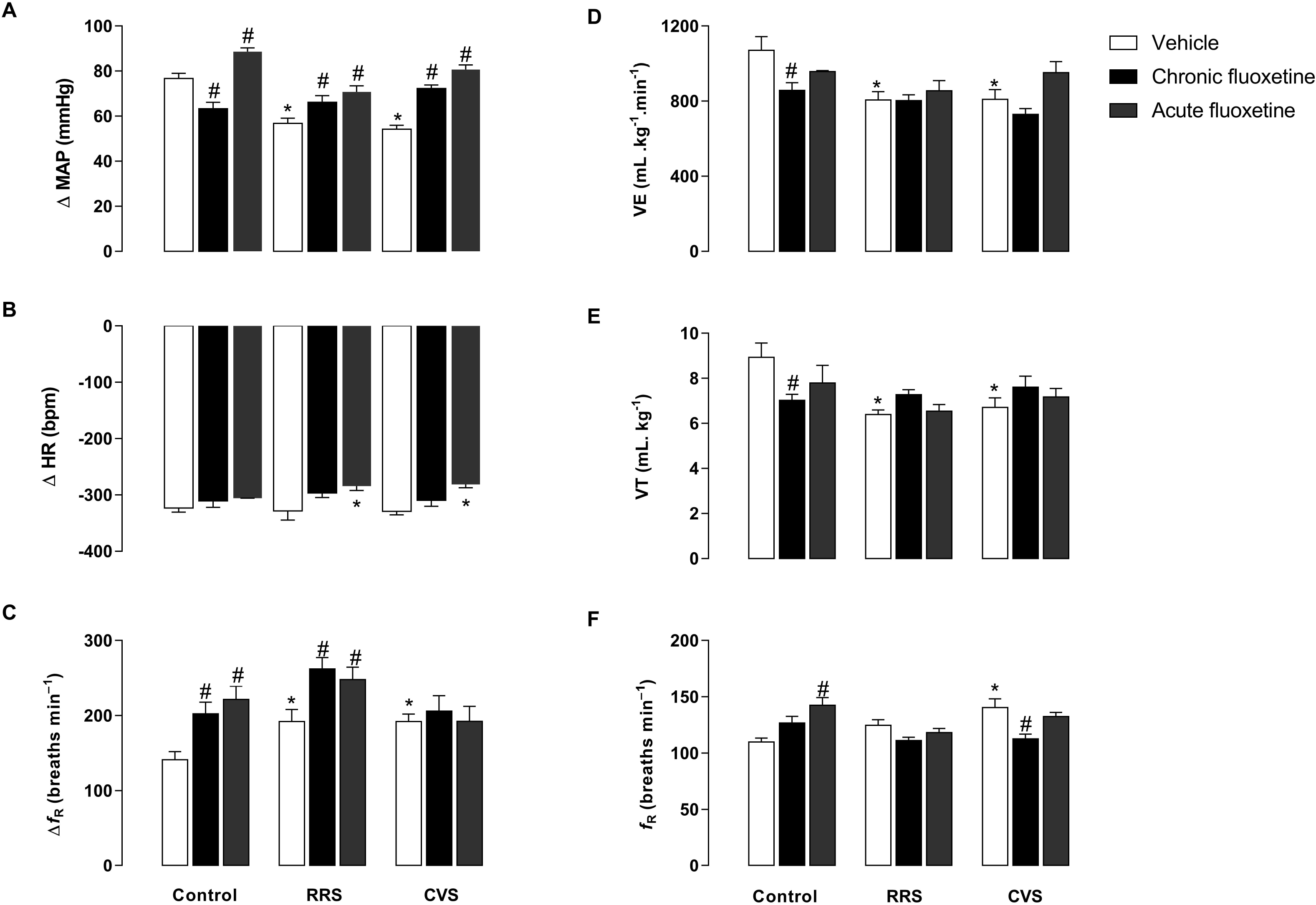

## References

Alboni, S., van Dijk, R.M., Poggini, S., Milior, G., Perrotta, M., Drenth, T., Brunello, N., Wolfer, D.P., Limatola, C., Amrein, I., Cirulli, F., Maggi, L., Branchi, I., 2017. Fluoxetine effects on molecular, cellular and behavioral endophenotypes of depression are driven by the living environment. Molecular psychiatry 22, 635.

Almeida, J., Duarte, J.O., Oliveira, L.A., Crestani, C.C., 2015. Effects of nitric oxide synthesis inhibitor or fluoxetine treatment on depression-like state and cardiovascular changes induced by chronic variable stress in rats. Stress 18, 462–474.

Alves, F.H., Crestani, C.C., Resstel, L.B., Correa, F.M., 2009. N-methyl-D-aspartate receptors in the insular cortex modulate baroreflex in unanesthetized rats. Auton Neurosci 147, 56–63.

Ampuero, E., Luarte, A., Santibanez, M., Varas-Godoy, M., Toledo, J., Diaz-Veliz, G., Cavada, G., Rubio, F.J., Wyneken, U., 2015. Two Chronic Stress Models Based on Movement Restriction in Rats Respond Selectively to Antidepressant Drugs: Aldolase C As a Potential Biomarker. The international journal of neuropsychopharmacology 18, pyv038.

Bahall, M., 2019. Prevalence and associations of depression among patients with cardiac diseases in a public health institute in Trinidad and Tobago. BMC psychiatry 19, 4.

Barres, C., Cheng, Y., Julien, C., 2004. Steady-state and dynamic responses of renal sympathetic nerve activity to air-jet stress in sinoaortic denervated rats. Hypertension 43, 629–635.

Bartlett, D., Tenney, S.M., 1970. Control of Breathing in Experimental Anemia. Resp Physiol 10, 384–&.

Berntson, G.G.C., J.T., 2007. Heart Rate Variability: Stress and Psychiatric Conditions. Dynamic Electrocardiography.

Carrive, P., 2006. Dual activation of cardiac sympathetic and parasympathetic components during conditioned fear to context in the rat. Clin Exp Pharmacol Physiol 33, 1251–1254.

Crestani, C.C., 2016. Emotional Stress and Cardiovascular Complications in Animal Models: A Review of the Influence of Stress Type. Frontiers in physiology 7, 251.

Crestani, C.C., Tavares, R.F., Guimaraes, F.S., Correa, F.M., Joca, S.R., Resstel, L.B., 2011. Chronic fluoxetine treatment alters cardiovascular functions in unanesthetized rats. Eur J Pharmacol 670, 527–533.

Cruz, F.C., Marin, M.T., Leao, R.M., Planeta, C.S., 2012. Behavioral and neuroendocrine effects of the exposure to chronic restraint or variable stress in early adolescent rats. Int J Dev Neurosci 30, 19–23.

Drorbaugh, J.E., Fenn, W.O., 1955. A barometric method for measuring ventilation in newborn infants. Pediatrics 16, 81–87.

Duarte, J.O., Planeta, C.S., Crestani, C.C., 2015. Immediate and long-term effects of psychological stress during adolescence in cardiovascular function: comparison of homotypic vs heterotypic stress regimens. Int J Dev Neurosci 40, 52–59.

Ferreira-Junior, N.C., Lagatta, D.C., Fabri, D.R., Alves, F.H., Correa, F.M., Resstel, L.B., 2017. Hippocampal subareas arranged in the dorsoventral axis modulate cardiac baroreflex function in a site-dependent manner in rats. Experimental physiology 102, 14–24.

Firmino, E.M.S., Kuntze, L.B., Lagatta, D.C., Dias, D.P.M., Resstel, L.B.M., 2019. Effect of chronic stress on cardiovascular and ventilatory responses activated by both chemoreflex and baroreflex in rats. The Journal of experimental biology 222.

Fitzgerald, R.S., 2000. Oxygen and carotid body chemotransduction: the cholinergic hypothesis - a brief history and new evaluation. Respir Physiol 120, 89–104.

Franchini, K.G., Krieger, E.M., 1993. Cardiovascular responses of conscious rats to carotid body chemoreceptor stimulation by intravenous KCN. J Auton Nerv Syst 42, 63–69.

Fuxe, K., 1965. Evidence for the Existence of Monoamine Neurons in the Central Nervous System. Iv. Distribution of Monoamine Nerve Terminals in the Central Nervous System. Acta Physiol Scand Suppl, SUPPL 247:237+.

Granjeiro, E.M., Scopinho, A.A., Correa, F.M., Resstel, L.B., 2011. Prelimbic but not infralimbic cortex is involved in the pressor response to chemoreflex activation in awake rats. Experimental physiology 96, 518–527.

Grippo, A.J., Beltz, T.G., Weiss, R.M., Johnson, A.K., 2006. The effects of chronic fluoxetine treatment on chronic mild stress-induced cardiovascular changes and anhedonia. Biol Psychiatry 59, 309–316.

Grippo, A.J., Moffitt, J.A., Johnson, A.K., 2002. Cardiovascular alterations and autonomic imbalance in an experimental model of depression. Am J Physiol Regul Integr Comp Physiol 282, R1333–1341.

Grippo, A.J., Moffitt, J.A., Johnson, A.K., 2008. Evaluation of baroreceptor reflex function in the chronic mild stress rodent model of depression. Psychosomatic medicine 70, 435–443.

Hodges, M.R., Opansky, C., Qian, B., Davis, S., Bonis, J., Bastasic, J., Leekley, T., Pan, L.G., Forster, H.V., 2004. Transient attenuation of CO2 sensitivity after neurotoxic lesions in the medullary raphe area of awake goats. J Appl Physiol (1985) 97, 2236–2247.

Hodges, M.R., Richerson, G.B., 2008. Contributions of 5-HT neurons to respiratory control: neuromodulatory and trophic effects. Respir Physiol Neurobiol 164, 222–232.

Holtman, J.R., Jr., 1988. Immunohistochemical localization of serotonin- and substance P-containing fibers around respiratory muscle motoneurons in the nucleus ambiguus of the cat. Neuroscience 26, 169–178.

Hong, L.Z., Huang, K.F., Hung, S.W., Kuo, L.T., 2017. Chronic fluoxetine treatment enhances sympathetic activities associated with abnormality of baroreflex function in conscious normal rats. European journal of pharmacology 811, 164–170.

Irvine, R.J., White, J., Chan, R., 1997. The influence of restraint on blood pressure in the rat. J Pharmacol Toxicol Methods 38, 157–162.

Ivanovs, R., Kivite, A., Ziedonis, D., Mintale, I., Vrublevska, J., Rancans, E., 2018. Association of depression and anxiety with cardiovascular co-morbidity in a primary care population in Latvia: a cross-sectional study. BMC public health 18, 328.

Jakubovski, E., Varigonda, A.L., Freemantle, N., Taylor, M.J., Bloch, M.H., 2016. Systematic Review and Meta-Analysis: Dose-Response Relationship of Selective Serotonin Reuptake Inhibitors in Major Depressive Disorder. The American journal of psychiatry 173, 174–183.

Kung, L.H., Scrogin, K.E., 2011. Serotonin nerve terminals in the dorsomedial medulla facilitate sympathetic and ventilatory responses to hemorrhage and peripheral chemoreflex activation. Am J Physiol Regul Integr Comp Physiol 301, R1367–1379.

Kuntze, L.B., Ferreira-Junior, N.C., Lagatta, D.C., Resstel, L.B., 2016. Ventral hippocampus modulates bradycardic response to peripheral chemoreflex activation in awake rats. Experimental physiology 101, 482–493.

Lagatta, D.C., Ferreira-Junior, N.C., Resstel, L.B., 2015. Medial prefrontal cortex TRPV1 channels modulate the baroreflex cardiac activity in rats. British journal of pharmacology 172, 5377–5389.

Lein, E.S., Hawrylycz, M.J., Ao, N., Ayres, M., Bensinger, A., Bernard, A., Boe, A.F., Boguski, M.S., Brockway, K.S., Byrnes, E.J., Chen, L., Chen, T.M., Chin, M.C., Chong, J., Crook, B.E., Czaplinska, A., Dang, C.N., Datta, S., Dee, N.R., Desaki, A.L., Desta, T., Diep, E., Dolbeare, T.A., Donelan, M.J., Dong, H.W., Dougherty, J.G., Duncan, B.J., Ebbert, A.J., Eichele, G., Estin, L.K., Faber, C., Facer, A., Fields, R., Fischer, S.R., Fliss, T.P., Frensley, C., Gates, S.N., Glattfelder, K.J., Halverson, K.R., Hart, M.R., Hohmann, J.G., Howell, M.P., Jeung, D.P., Johnson, R.A., Karr, P.T., Kawal, R., Kidney, J.M., Knapik, R.H., Kuan, C.L., Lake, J.H., Laramee, A.R., Larsen, K.D., Lau, C., Lemon, T.A., Liang, A.J., Liu, Y., Luong, L.T., Michaels, J., Morgan, J.J., Morgan, R.J., Mortrud, M.T., Mosqueda, N.F., Ng, L.L., Ng, R., Orta, G.J., Overly, C.C., Pak, T.H., Parry, S.E., Pathak, S.D., Pearson, O.C., Puchalski, R.B., Riley, Z.L., Rockett, H.R., Rowland, S.A., Royall, J.J., Ruiz, M.J., Sarno, N.R., Schaffnit, K., Shapovalova, N.V., Sivisay, T., Slaughterbeck, C.R., Smith, S.C., Smith, K.A., Smith, B.I., Sodt, A.J., Stewart, N.N., Stumpf, K.R., Sunkin, S.M., Sutram, M., Tam, A., Teemer, C.D., Thaller, C., Thompson, C.L., Varnam, L.R., Visel, A., Whitlock, R.M., Wohnoutka, P.E., Wolkey, C.K., Wong, V.Y., Wood, M., Yaylaoglu, M.B., Young, R.C., Youngstrom, B.L., Yuan, X.F., Zhang, B., Zwingman, T.A., Jones, A.R., 2007. Genome-wide atlas of gene expression in the adult mouse brain. Nature 445, 168–176.

Li, Y.Q., Takada, M., Mizuno, N., 1993. The sites of origin of serotoninergic afferent fibers in the trigeminal motor, facial, and hypoglossal nuclei in the rat. Neurosci Res 17, 307–313.

Lino-de-Oliveira, C., Sales, A.J., Del Bel, E.A., Silveira, M.C., Guimaraes, F.S., 2001. Effects of acute and chronic fluoxetine treatments on restraint stress-induced Fos expression. Brain Res Bull 55, 747–754.

Low, C.A., Thurston, R.C., Matthews, K.A., 2010. Psychosocial factors in the development of heart disease in women: current research and future directions. Psychosom Med 72, 842–854.

Lu, Y., Ho, C.S., Liu, X., Chua, A.N., Wang, W., McIntyre, R.S., Ho, R.C., 2017. Chronic administration of fluoxetine and pro-inflammatory cytokine change in a rat model of depression. PloS one 12, e0186700.

Marin, M.T., Cruz, F.C., Planeta, C.S., 2007. Chronic restraint or variable stresses differently affect the behavior, corticosterone secretion and body weight in rats. Physiology & behavior 90, 29–35.

McFarlane, A., Kamath, M.V., Fallen, E.L., Malcolm, V., Cherian, F., Norman, G., 2001. Effect of sertraline on the recovery rate of cardiac autonomic function in depressed patients after acute myocardial infarction. American heart journal 142, 617–623.

Nakata, T., Berard, W., Kogosov, E., Alexander, N., 1993. Cardiovascular change and hypothalamic norepinephrine release in response to environmental stress. Am J Physiol 264, R784–789.

Pacher, P., Kecskemeti, V., 2004. Cardiovascular side effects of new antidepressants and antipsychotics: new drugs, old concerns? Curr Pharm Des 10, 2463–2475.

Papademetriou, V., Doumas, M., Faselis, C., Tsioufis, C., Douma, S., Gkaliagkousi, E., Zamboulis, C., 2011. Carotid baroreceptor stimulation for the treatment of resistant hypertension. International journal of hypertension 2011, 964394.

Paton, J.F., Deuchars, J., Li, Y.W., Kasparov, S., 2001. Properties of solitary tract neurones responding to peripheral arterial chemoreceptors. Neuroscience 105, 231–248.

Resstel, L.B., Fernandes, K.B., Correa, F.M., 2004. Medial prefrontal cortex modulation of the baroreflex parasympathetic component in the rat. Brain Res 1015, 136–144.

Resstel, L.B., Joca, S.R., Guimaraes, F.G., Correa, F.M., 2006. Involvement of medial prefrontal cortex neurons in behavioral and cardiovascular responses to contextual fear conditioning. Neuroscience 143, 377–385.

Roche, M., Harkin, A., Kelly, J.P., 2007. Chronic fluoxetine treatment attenuates stressor-induced changes in temperature, heart rate, and neuronal activation in the olfactory bulbectomized rat. Neuropsychopharmacology 32, 1312–1320.

Roose, S.P., Glassman, A.H., Attia, E., Woodring, S., Giardina, E.G., Bigger, J.T., Jr., 1998. Cardiovascular effects of fluoxetine in depressed patients with heart disease. Am J Psychiatry 155, 660–665.

Tavares, R.F., Correa, F.M., 2006. Role of the medial prefrontal cortex in cardiovascular responses to acute restraint in rats. Neuroscience 143, 231–240.

Taylor, N.C., Li, A., Green, A., Kinney, H.C., Nattie, E.E., 2004. Chronic fluoxetine microdialysis into the medullary raphe nuclei of the rat, but not systemic administration, increases the ventilatory response to CO2. J Appl Physiol (1985) 97, 1763–1773.

Taylor, N.C., Li, A., Nattie, E.E., 2006. Ventilatory effects of muscimol microdialysis into the rostral medullary raphe region of conscious rats. Respir Physiol Neurobiol 153, 203–216.

Thor, K.B., Helke, C.J., 1987. Serotonin- and substance P-containing projections to the nucleus tractus solitarii of the rat. J Comp Neurol 265, 275–293.

Weissheimer, K.V., Machado, B.H., 2007. Inhibitory modulation of chemoreflex bradycardia by stimulation of the nucleus raphe obscurus is mediated by 5-HT3 receptors in the NTS of awake rats. Autonomic Neuroscience-Basic & Clinical 132, 27–36.

Willner, P., 1990. Animal models of depression: an overview. Pharmacol Ther 45, 425–455.

Willner, P., 1997. Validity, reliability and utility of the chronic mild stress model of depression: a 10-year review and evaluation. Psychopharmacology (Berl) 134, 319–329.

Willner, P., 2005. Chronic mild stress (CMS) revisited: consistency and behavioural-neurobiological concordance in the effects of CMS. Neuropsychobiology 52, 90–110.

Willner, P., Muscat, R., Papp, M., 1992. Chronic Mild Stress-Induced Anhedonia - a Realistic Animal-Model of Depression. Neurosci Biobehav R 16, 525–534.

